# FoldVision: A compute-efficient atom-level 3D protein encoder

**DOI:** 10.64898/2026.01.23.701326

**Authors:** Alexander Kroll, Shantanu Yadav, Martin J. Lercher

## Abstract

Protein function emerges from three-dimensional structure, yet many large-scale protein prediction pipelines still rely solely on linear sequence embeddings. Although multiple structure-aware protein networks have been proposed, they often omit atom-level details and struggle to capture the detailed chemistry of binding sites. Here, we introduce FoldVision, a compute-efficient 3D convolutional neural network that voxelizes every heavy atom, learns rotation-robust representations, and is pre-trained on over 500 000 AlphaFold-2 structures, which is more than two orders of magnitude less data than used to train modern protein language models. Despite its compact size of 123 million parameters, FoldVision outperforms or matches state-of-the-art protein encoders on four benchmarks that require fine structural resolution: enzyme–substrate classification, transporter–substrate classification, drug–kinase inhibition, and drug–target inhibition prediction. A simple ensemble with a sequence-based model consistently improves performance across all benchmarks beyond any individual model. This indicates that FoldVision learns structural signals that are complementary to those extracted by sequence-based models. This study demonstrates that full-atom protein 3D CNNs are both tractable and superior to protein language models alone for structure-dependent tasks.

## Introduction

Protein sequence databases are expanding orders of magnitude faster than experimental annotation can keep pace, widening the knowledge gap between sequence and function [1]. Scalable *in silico* methods are therefore needed to translate raw sequences into biological insights.

Today’s leading approaches encode sequences with transformer-based protein language models (PLMs) such as ESM-2 and ProtT5 [2, 3]. Trained on hundreds of millions of sequences, PLMs capture evolutionarily conserved motifs and long-range residue dependencies that underpin function. Function, however, is ultimately determined by the three-dimensional protein geometry. While large PLMs can infer structure implicitly [2, 4], many applications demand atom-level accuracy. Supplying models with explicit coordinates obtained from experiments or structure prediction models should therefore benefit tasks that require precise active- or binding-site geometries.

Recent work has begun to feed deep learning models with explicit structural data. Approaches such as DeepFRI [5], GearNet [6], and ESM-GearNet [7] represent proteins as graphs, where residues are treated as nodes encoded by PLM embeddings and edges capture covalent bonds or spatial proximity. The result is a protein graph that can be processed by a graph neural network (GNN) [8].

These protein graphs capture residue connectivity but ignore the precise arrangement of atoms. Bond angles, side-chain conformations, and binding pocket geometries are either oversimplified into edge features or omitted altogether. Because large PLMs already infer many residue–residue contacts from sequence alone, augmenting them with a connectivity graph yields only incremental gains, such as faster convergence and small accuracy improvements. For instance, in the extensive ESM-GearNet study, most investigated protein graph encodings only slightly outperformed the sequence-only ESM-2 baseline, with some even performing worse [7].

ESM-IF [9] and ProstT5 [10] are structure-aware protein transformers trained to translate between structural and sequence tokens. While these models move closer to explicitly encoding protein structure, they discretize the protein and omit full atom-level detail. For tasks that depend on subtle geometric features, such as binding-partner complementarity, electrostatics, or active-site geometry, this abstraction can reduce predictive performance. In contrast, a model that operates directly on the full, continuous 3D coordinates of all atoms should be better able to capture the underlying physics and chemistry that govern molecular interactions.

Convolutional neural networks (CNNs), originally popularized in computer vision for 2D image classification and object detection, have been adapted to volumetric 3D data [11]. Between 2017 and 2020, three-dimensional CNNs (3D CNNs) were popular choices for processing 3D protein representations. Early studies used local or simplified input representations: DeepSite locates binding pockets from local grids [12], EnzyNet predicts EC numbers from the atomic positions of the protein carbon backbone [13], and VoroCNN converts atomic coordinates into a contact graph with Voronoi diagrams to evaluate how well each region of a structure is folded [14]. 3DCNN-MQA extends these ideas and uses full atomic 3D CNNs to assess the quality of predicted protein structures [15]. All these methods were constrained by limited GPU memory and the absence of today’s large public structure datasets such as AlphaFoldDB [16], resulting in models with simplified or localized representations without large-scale pretraining.

Despite recent hardware and data gains, deep learning models acting on full atomic protein 3D structures remain uncommon. It is believed that applying 3D CNNs to entire, high-resolution protein structures is computationally inefficient due to the sparsity of protein atoms in 3D space and the significant memory requirements [5, 17]. Additionally, proteins lack a canonical orientation, whereas vanilla CNNs are not rotation and translation invariant. Solutions therefore require either invariant architectures or extensive orientation augmentation. Nevertheless, recent evidence shows that 3D CNNs for proteins can add complementary information beyond what sequence-based models learn, demonstrated by successfully predicting masked amino acids from local 3D environments [18].

Here we introduce FoldVision, a 3D CNN encoder that processes full-atom protein structures (Figure 1). The model maps each atom to a 3D voxel grid and is pre-trained on ∼ 500 000 structures from the AlphaFoldDB [16]. We aimed to achieve rotation-robust predictions through extensive 3D augmentations, while grid sizes adaptive to protein size mitigate memory wastage for smaller proteins. Because the pipeline uses only predicted structures, it can be applied to the vast protein sequence space without requiring experimental data.

**Figure 1.**
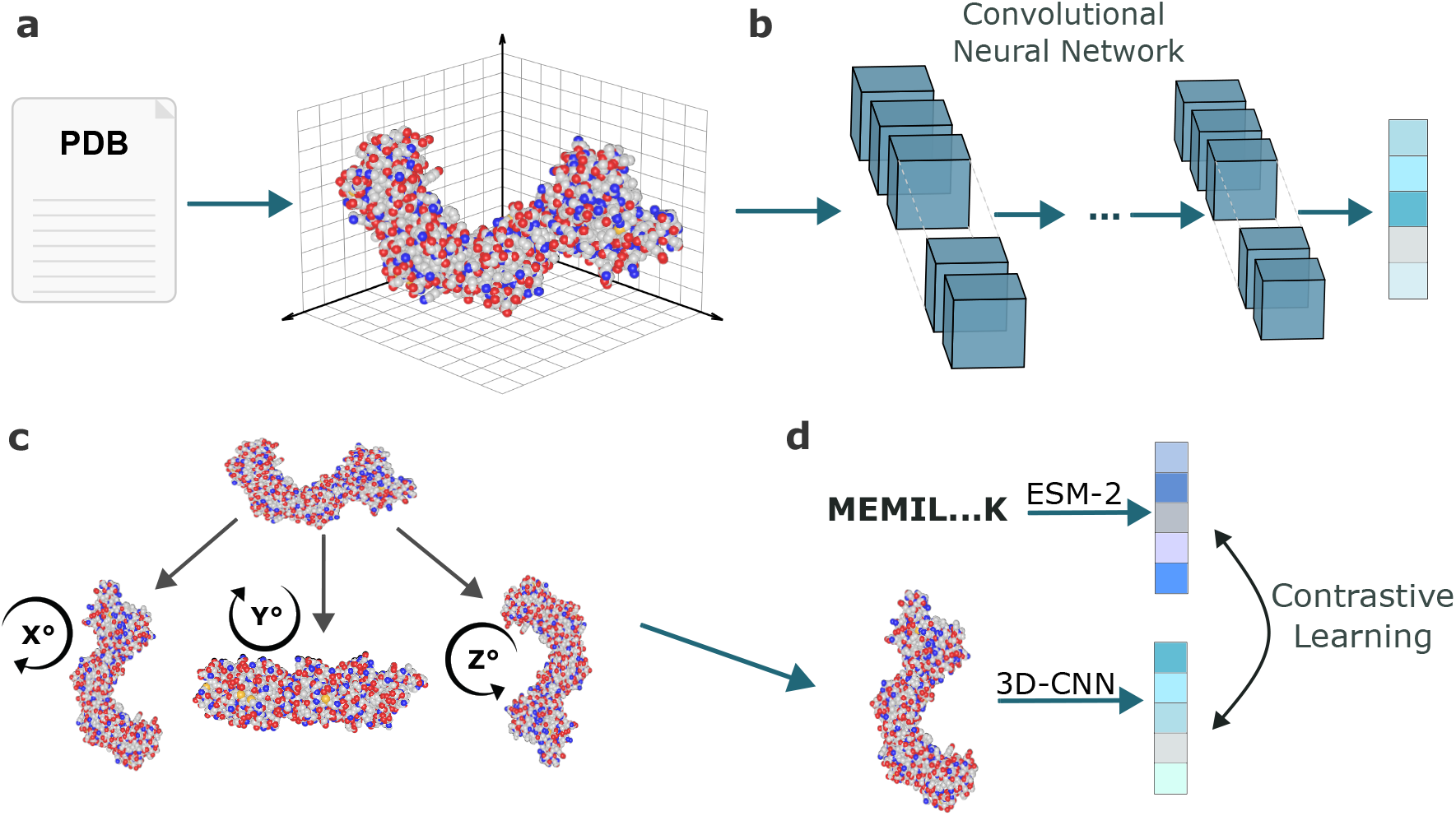
Overview of the FoldVision workflow. **(a)** PDB structures are voxelized into 3D grids that encode the positions of the most common heavy atoms in the protein. **(b)** The resulting grid serves as input to a 3D convolutional neural network (3D CNN). Convolutional features are globally averaged to yield a fixed-length protein embedding. **(c)** To enforce rotational invariance, each structure undergoes random rotations during training. **(d)** FoldVision is pre-trained with a contrastive objective that pulls embeddings toward the corresponding ESM-2 embedding of the same protein and pushes them away from those of other proteins.

The goal of this study was not to introduce an entirely new architectural paradigm, but to demonstrate that full-atom 3D CNNs, when trained at scale with modern compute and comprehensive structural data, achieve state-of-the-art performance. We show that ensembling FoldVision with a PLM surpasses current best methods for tasks that depend on fine structural details, such as transporter and enzyme substrate prediction, as well as drug-target interaction prediction. This performance is achieved while requiring at least 5 times fewer learnable parameters and more than two orders less pre-training data compared to modern protein language models.

## Results

### Converting protein structures into 3D voxel grids

FoldVision, our 3D CNN protein encoder, encodes full-atom protein structures as five-channel 3D voxel grids at 1.0 Å resolution (Figure 1). Each channel stores the local density of one of the five most common heavy-atom types: carbon, nitrogen, oxygen, sulfur, and phosphorus. For each voxel these values are stored in a 5-dimensional vector, and hence each protein is represented by a 4D array: three spatial dimensions plus one channel dimension.

Instead of one-hot encoding the presence of atoms at voxels, each atom contributes a continuous density to the 3D grid. For an atom of type *c* at position 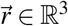, its contribution to the *c*-channel of the voxel centered at 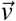 is 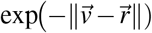. In contrast to binary encodings, this reduces discretizations artifacts, preserves local geometry, and creates a smoother, more continuous input.

To process proteins of widely differing sizes while limiting empty space in the input boxes, we embed each structure in the smallest of four cubic boxes (64^3^, 96^3^, 128^3^, or 160^3^ Å ^3^) that fully encloses all atoms. This adaptive boxing minimizes the number of empty voxels and thus lowers memory and compute requirements. Structures that exceed the 160^3^-voxel grid are cropped at the bounding box.

Because standard 3D CNNs are orientation-dependent, FoldVision produces distinct embeddings for rotated copies of the same structure. To this end, every time a protein is presented to the model, we rotate it by a random degree around one of the three cartesian axes or leave it unchanged (Figure 1c); each of the four options is chosen with equal probability. This input augmentation enables the network to learn more robust embeddings and predictions, which are less affected by arbitrary rotations of the input protein.

### FoldVision utilizes 5 blocks of 3D convolutional operations

FoldVision consists of a three-dimensional convolutional neural network (3D CNN) that operates directly on the atomic-density voxel grid described above. Each protein input box is processed by five FoldVision blocks (Table S1), where each block contains one or two 3D convolution layers, a group normalization layer, and a ReLU layer, followed by a 2 × 2 × 2 max-pooling operation that halves the spatial resolution. Kernel sizes for the convolutions in layers 1–5 are 7^3^, 5^3^, 3^3^, 3^3^, and 3^3^, respectively. After the final layer, the 3D feature map is reduced in resolution and each voxel is encoded as a 1 024-dimensional vector. Global average pooling over all voxels yields a fixed 1 024-dimensional representation for every protein, independent of the input box size.

For protein-small molecule interaction prediction tasks, this protein vector is concatenated with a 768-dimensional MoLFormer-XL small-molecule embedding [19]. The resulting vector is fed to a two-layer feed-forward network (256 hidden units) with batch normalization, ReLU activation, and dropout to produce a task-specific output prediction. In total, FoldVision contains ∼ 123 M learnable parameters, which is much fewer than the number of parameters of current protein language models that range up to 15 billion [2, 3, 20].

### Contrastive structure–sequence pre-training

We pre-trained FoldVision on more than 500 000 AlphaFold-2-predicted protein structures using a contrastive alignment objective. Specifically, we projected ESM-2 sequence embeddings [2] and FoldVision structure embeddings into a shared space, and optimized a contrastive loss that pulls together matched sequence–structure embeddings from the same protein while pushing apart embeddings from different proteins (Figure 1d). This pre-training teaches FoldVision to map a protein’s 3D shape into the same “meaning space” learned by ESM-2 from sequences, allowing it to inherit rich biological information and providing a useful prior for downstream prediction tasks.

We also tested a multi-task regression pre-training: The model was trained to predict 23 *in silico* computed Rosetta scores (Table S2), which summarize per-residue and whole-protein physicochemical properties. However, this resulted in worse downstream prediction performance than the contrastive pretraining (see subsection “Multi-view ensembling is important for obtaining high model performance”). Consequently, all task-specific models are initialized with the contrastively pre-trained weights; the task-specific prediction head is replaced with a freshly initialized fully connected neural network, and the entire network is then fine-tuned end-to-end.

### Experimental setup for a fair comparison with other protein encoders

We compared FoldVision against sequence-based protein language models (ProtT5 [3], ESM-2_150M_ with 150M parameters, and ESM-2_650M_ with 650M parameters [2]), structure-encoding transformers (ProstT5 [10] and ESM-IF [9]), and the 3D convolutional model 3DCNN-MQA [15].

For each task, all models were fine-tuned with the same protocol: we tuned the learning rate, trained until validation performance stopped improving, and kept the best checkpoint based on the validation set. We then evaluated the models on independent test sets. For the protein-small molecule prediction tasks, we used for all models fixed MoLFormer-XL embeddings to encode the small molecule [19]. On top of the extracted molecule and protein embeddings, we used the same two-layer prediction head as in FoldVision (256 hidden units, batch normalization, ReLU).

The fine-tuning pipeline is deliberately simple and likely underestimates for all models the peak accuracy achievable with a fully optimized pipeline. For example, fine-tuning the small-molecule encoder or feeding the fine-tuned protein embeddings as input for separate machine learning models can yield higher scores [21]. However, we chose the identical, simple pipeline for all models to isolate the effect of the protein encoder.

### FoldVision accurately predicts interactions between small molecule drugs and target proteins

To quantify the practical benefit of FoldVision’s atom-level information, we benchmarked it against other protein encoders on four problems that require accurate pocket recognition and ligand matching: (i) prediction of the inhibition constant IC_50_ for drug–target pairs using the BindingDB dataset [22], (ii) prediction of the dissociation constants *K*_*d*_ for pairs of drugs and protein kinases using the Davis dataset [23], (iii) predicting whether a small molecule is a substrate of a given enzyme [21], and (iv) predicting whether a small molecule is a substrate of a given transporter [24]. For the Davis dataset, we employed cold target splitting, where the proteins in the test and validation sets do not overlap with those in the training set. For details on the datasets, see Table S3.

Across both drug–target interaction prediction tasks, FoldVision provides complementary signal that improves predictive performance (Table 1). On the BindingDB dataset, FoldVision outperforms all other models. Using a one-sided Wilcoxon signed-rank test, we compared FoldVision’s absolute prediction errors with those of each other protein-encoding model and found the improvements to be statistically significant (*p* < 10^−282^). Notably, almost all other protein encoders were pre-trained on hundreds of millions of proteins, whereas FoldVision was pre-trained on only 0.5 million structures.

**Table 1.** Prediction performance of sequence- and structure-based models on the BindingDB (IC50) and Davis (cold target) test sets. Arrows show whether higher (↑) or lower (↓) values indicate better performance. Bold numbers highlight the best performance for each metric. *R*^2^: Coefficient of determination; MAE: Mean absolute error.

| Model | Input | BindingDB (IC50) |  |  | Davis (cold target) |  |  |
| --- | --- | --- | --- | --- | --- | --- | --- |
| | | MAE ↓ | $R^2$ ↑ | Pearson r ↑ | MAE ↓ | $R^2$ ↑ | Pearson r ↑ |
| ESM-2<sub>150M</sub> | Seq. | 0.705 | 0.602 | 0.780 | 0.401 | 0.623 | 0.793 |
| ESM-2<sub>650M</sub> | Seq. | 0.710 | 0.597 | 0.775 | 0.379 | 0.671 | 0.787 |
| ProtT5 | Seq. | 0.716 | 0.589 | 0.768 | 0.360 | 0.674 | 0.827 |
| ESM-IF | Struct. | 0.777 | 0.523 | 0.730 | 0.398 | 0.597 | 0.777 |
| 3DCNN-MQA | Struct. | 0.705 | 0.598 | 0.775 | 0.426 | 0.386 | 0.650 |
| FoldVision | Struct. | **0.627** | **0.687** | **0.830** | 0.356 | 0.645 | 0.804 |
| ESM-FoldVision | Seq.+Struct. | 0.643 | 0.667 | 0.818 | **0.348** | **0.686** | **0.835** |

On the Davis dataset, FoldVision outperforms all structure-based models and achieves performance comparable to the sequence-based models ESM-2_650M_ and ProtT5: FoldVision attains a better mean absolute error, while *R*^2^ and Pearson’s *r* are slightly worse. Averaging the predictions of FoldVision and ESM-2_650M_ yields an ensemble that further improves model performance on the Davis dataset. The ensemble’s absolute errors are significantly lower than those of each individual protein-encoding model (one-sided Wilcoxon signed-rank test, *p* < 10^−10^).

On both binary classification benchmarks, the transporter–substrate prediction (TSP) and enzyme–substrate prediction (ESP) tasks, FoldVision matches or outperforms the performance of all other protein encoding models, only ProtT5 performs slightly better on the TSP dataset (Table 2). Notably, the ESM–FoldVision ensemble outperforms every individual model on all metrics, indicating that the two models combine complementary strength: while ESM-2 likely captures evolutionary sequence motifs, FoldVision contributes explicit 3D pocket geometry. For both prediction tasks, we performed a McNemar’s test to assess whether there are significant differences in the error rates of the ESM-FoldVision ensemble and each model alone, and the differences were found to be significant (*p* < 10^−3^ for TSP and *p* < 10^−6^ for ESP).

**Table 2.** Prediction performance of ESM-2 and FoldVision on the transporter-substrate (TSP) and enzyme-substrate (ESP) test sets. Higher values indicate better performance. Bold numbers highlight the best performance for each metric. Acc: Accuracy; ROC-AUC: Receiver Operating Characteristic – Area Under the Curve; MCC: Matthews correlation coefficient; Prec: Precision.

| Model | Input | Transporter-Substrate (TSP) |  |  |  | Enzyme-Substrate (ESP) |  |  |  |
| --- | --- | --- | --- | --- | --- | --- | --- | --- | --- |
|  |  | Acc | ROC-AUC | MCC | Prec | Acc | ROC-AUC | MCC | Prec |
| ESM-2<sub>150M</sub> | Seq. | 88.5% | 0.943 | 0.725 | 0.771 | 91.8% | 0.957 | 0.789 | 0.860 |
| ESM-2<sub>650M</sub> | Seq. | 89.3% | 0.947 | 0.743 | 0.789 | 91.9% | 0.955 | 0.791 | 0.863 |
| ProtT5 | Seq. | 91.8% | 0.964 | 0.801 | 0.835 | 91.5% | 0.951 | 0.782 | 0.858 |
| ProstT5 | Struct. | 83.8% | 0.914 | 0.622 | 0.683 | 87.6% | 0.929 | 0.663 | 0.719 |
| ESM-IF | Struct. | 89.3% | 0.947 | 0.738 | 0.801 | 91.3% | 0.953 | 0.776 | 0.848 |
| 3DCNN-MQA | Struct. | 84.6% | 0.902 | 0.628 | 0.711 | 81.2% | 0.864 | 0.536 | 0.636 |
| FoldVision | Struct. | 91.2% | 0.955 | 0.779 | 0.871 | 91.8% | 0.951 | 0.786 | 0.889 |
| ESM-FoldVision | Seq.+Struct. | **92.4%** | **0.966** | **0.817** | **0.876** | **92.9%** | **0.961** | **0.816** | **0.903** |

### Structure-based models are less effective for homology-reliant prediction tasks

We hypothesize that the relative value of sequence and structure information depends on the task: sequence- and homology-based information is likely more relevant for broad classification problems, whereas detailed affinity prediction requires high-resolution structural context. To also test FoldVision’s utility on problems believed to be well explained by sequence homology, we evaluated its performance on Gene Ontology (GO) annotation [25] predictions, which describe molecular functions, biological processes, and cellular components of proteins.

GO inference is often approached by nearest-neighbor searches: transferring annotations from the closest BLAST hit already achieves reasonable results in the CAFA GO Term prediction challenge [26, 27]. Consequently, a sequence-based transformer with global attention might outperform a structure-aware CNN that relies on local 3D filters.

We fine-tuned FoldVision and ESM-2_650M_ for predicting molecular-function GO terms and evaluated them on an independent test set. FoldVision reached an *F*_max_ of 0.57, below ESM-2_650M_’s 0.62. Averaging their prediction raised performance marginally to 0.63. Thus, when functional inference depends mainly on conserved motifs or overall homology, structure encoders indeed add limited or no value.

### Prediction accuracy varies with sequence and structural similarity

To examine under which conditions sequence and structure encoder perform best, we partitioned the transporter–substrate test set according to three criteria and compared FoldVision to ESM-2_650M_ (Figure 2 a-c). We first binned proteins by their maximum pairwise sequence identity compared to the training proteins (Figure 2a). In high-identity bins, FoldVision outperforms ESM-2. This indicates that the functional impact of a few amino acid substitutions is captured more effectively from 3D structures than from linear sequences. As identity drops, the gap narrows. In the lowest-identity bin, the models perform similarly, but averaging their predictions improves accuracy compared to each model alone.

**Figure 2.**
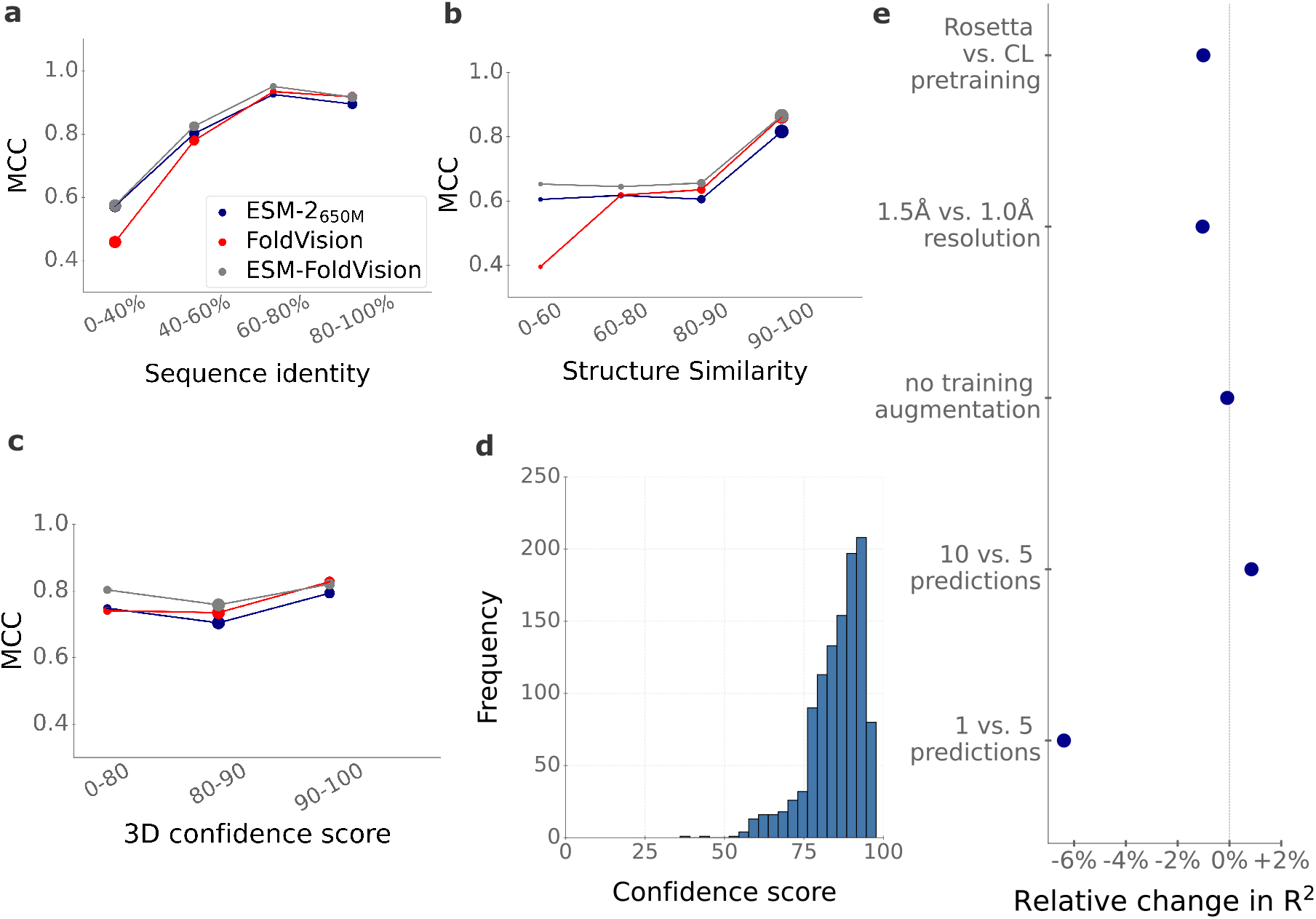
The ESM–FoldVision ensemble combines complementary signals from sequence and structure. Matthews correlation coefficient (MCC) on different subsets of the transporter–substrate test set based on (a) maximum sequence identity to the training set and (b) maximum TM-score (global structural similarity) to any training structure. (c) MCC binned by AlphaFold-2 average pLDDT confidence score of the predicted structures. The areas of the circles in panels (a) - (c) are proportional to the number of data points in each subset. (d) Distribution of pLDDT scores across test proteins. (e) Relative change in IC50 regression performance for alternative architectural or training choices, expressed with respect to the selected FoldVision design.

We repeated the analysis using the maximum pairwise TM-score, a measure of global structural similarity [28], instead of sequence identity (Figure 2b). The pattern is similar: FoldVision performs best when the test fold is well-represented in the training set, whereas ESM-2 performs better when similar structures are absent from the training set. Across all similarity ranges, the ESM–FoldVision ensemble outperforms both single models, showing their complementarity.

FoldVision relies on AlphaFold-2–predicted structures. We examined how prediction accuracy varies with the quality of those structures [16], estimated as the mean per-residue confidence score (pLDDT; Figure 2c). Stratifying the test set by pLDDT shows that FoldVision performs best on high-confidence structures (pLDDT > 90), suggesting that experimental structures could further improve accuracy. On low-confidence models, FoldVision performs comparably to ESM-2. Most test proteins, however, fall into the high-confidence range (Figure 2d), indicating that the observed gains reflect the majority of cases rather than a small subset.

### Multi-view ensembling is important for obtaining high model performance

We quantified the effect of key architectural and training choices by starting from the reference FoldVision model used for the IC_50_ regression benchmark and perturbing one component at a time (Figure 2e). Reported changes are relative to the reference coefficient of determination *R*^2^ = 0.687 achieved with the full model.

During inference, FoldVision averages predictions from five randomly augmented views of each protein. Reducing the ensemble to a single view lowered *R*^2^ by 6.4%, confirming that multiple orientations are essential for robust inference. Expanding to ten instead of five views yields a small 0.9% gain. Strikingly, disabling rotational augmentations during fine-tuning had virtually no impact (–0.1 %), implying that the network learns a stable representation already during pre-training and can be fine-tuned on unaugmented data.

Replacing the contrastive alignment to ESM-2 embeddings with multi-task regression on 23 Rosetta energy terms decreased *R*^2^ by 1.0%. Both objectives therefore produce competitive encoders, with the contrastive objective offering a slight advantage for affinity prediction. Voxel size resolution also influences performance: reducing the resolution from 1.0Å to 1.5Å per voxel reduced *R*^2^ by 1.1%. This indicates that atom-level resolution is important but can be modestly relaxed without catastrophic loss.

### 3D CNNs are compute- and data-efficient but storage-heavy

To assess the requirements for running sequence-based transformer models and 3D structure–based CNNs, we compared compute, data, and storage requirements for FoldVision and the ESM-2 models (Figure 3). A single forward pass requires on average 415 Giga Floating Point Operations (GFLOPs) for FoldVision, 520 GFLOPs for ESM-2_650M_, and 140 GFLOPs for ESM-2_150M_ (Figure 3b). Despite similar numbers of mathematical operations compared to ESM-2_650M_, FoldVision processes 1 000 proteins on one A100 GPU in 20 seconds during training, versus 215 seconds for ESM-2_150M_ and 426 seconds for the ESM-2_650M_ (Figure 3c). This suggests that computations for the FoldVision CNN with only five FoldVision blocks can be parallelized more efficiently than those required for the ESM-2 models that have many more encoder layers. On the transporter-substrate prediction task, FoldVision achieved its best performance after 52 epochs and ESM-2_650M_ after 47 epochs, indicating that faster inference does not come at the cost of slower convergence.

Pre-training requirements diverge even more: ESM-2_650M_ ran for ∼ 7 days on 128 H100 GPUs and 250 M different protein sequences, whereas FoldVision finished pre-training in 48 hours on four A100 GPUs using structures from ∼ 0.5 M proteins. The FoldVision pre-training thus used only about 1% of the GPU-hours. The trade-off is disk space: For example, the 5 800 proteins from the transporter dataset require only 3 MB when storing the linear amino acid sequence (in FASTA format) but ∼ 75 GB as float32 coordinate NumPy arrays (Figure 3d). Thus, FoldVision is cheaper in compute and data, but requires larger storage for 3D inputs.

**Figure 3.**
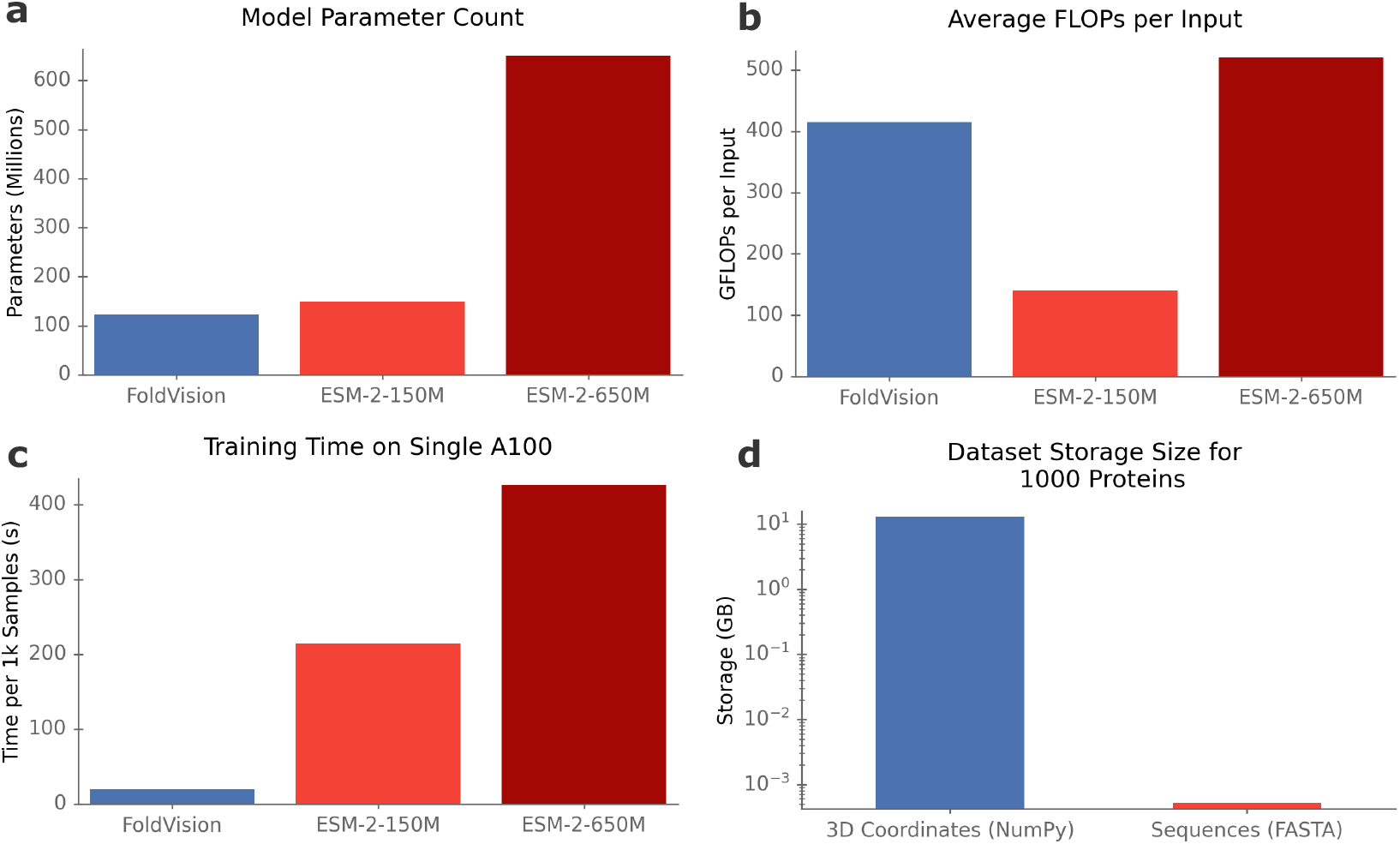
FoldVision is compute- and runtime-efficient but storage-intensive. Comparison of FoldVision with ESM–2_150M_ and ESM–2_650M_ for **(a)** trainable parameters, **(b)** average GFLOPs per protein, **(c)** training time to process 1 000 proteins (A100 GPU), and **(d)** disk space required to store input data for 1 000 proteins.

## Discussion

FoldVision aims to convert the vast amount of available protein structures into detailed biological insights and applies 3D CNNs directly to high-resolution protein structures. By explicitly using atom-level information, the model adds important information in prediction tasks that require accurate representations of the precise geometry of a protein’s structure. Despite being pre-trained on orders of magnitude fewer proteins and computational resources, FoldVision outperforms or matches the performance of widely used sequence-based models as well as other structure-based models on four benchmarks of protein-small molecule interaction prediction tasks.

While prior work has encoded full-atom protein structures with 3D CNNs [15], we are, to our knowledge, the first to develop a general-purpose, large-scale full-atom 3D protein encoder that takes advantage of modern GPUs and rapidly expanding structural databases. Our aim was to provide a proof-of-concept that large-scale pretraining and fine-tuning of full-atom structure encoders is computationally feasible, complements sequence-based transformers, and can outperform sequence-only standards such as ESM-2. We approached this goal using standard 3D CNNs as a straightforward baseline. However, rotation-invariant CNNs [29] and point-cloud models [30] may be even better suited.

FoldVision achieves high predictive performance without annotations of functionally relevant structural regions, such as binding sites, indicating that it can infer and encode such regions autonomously. Adapting visualization methods such as Grad-CAM [31] to 3D convolutions could reveal the voxels that drive individual predictions. This could pinpoint candidate binding pockets for downstream analysis or experimental validation [32].

Effective pre-training strongly influences downstream performance of protein encoders. Here, we compared two pre-training objectives on approximately 500 000 structures: (i) Regression on 23 Rosetta scores, reflecting physicochemical properties and (ii) contrastive learning against ESM-2 embeddings of the linear amino acid sequence. The computer vision literature offers a rich catalog of alternative objectives, such as masked voxel prediction, jigsaw tasks, rotation prediction, and multi-view contrastive learning, which could be transferred to volumetric protein data and could yield even more expressive representations. Furthermore, the depth and breadth of FoldVision’s neural network are modest compared to large 2D-CNNs that follow well-established scaling laws [33]. It is not yet known whether similar laws apply in the sparse, high-resolution regime of protein space, but systematically exploring larger, deeper 3D CNNs is a promising path toward further improvements.

A current limitation is that FoldVision encodes only a single rigid snapshot of each structure. Proteins, however, are dynamic and sample ensembles of conformations that govern function and binding partner affinity. Incorporating alternative snapshots, such as those generated by DynaMine [34], C-Fold [35], MD-Gen [36], AF2*χ* [37], or molecular-dynamics simulations [38], would allow the network to observe how pockets expand or contract. As demonstrated here, even simple augmentations such as random rotations improve performance; dynamic augmentations should supply even richer geometric information and yield more robust predictions.

FoldVision’s strong performance on structure-sensitive tasks suggests several new applications. On the one hand, predicting the functional impact of mutations could profit from the detailed encoding of the geometry of all atoms; accurate mutant structures can be produced by tools such as PDBFixer [39]. Furthermore, protein–protein interaction predictions depend on recognizing complementary surface patches across two macromolecules, an inherently structural problem well aligned with FoldVision’s capabilities.

## Methods

### Software and code availability

All models are implemented in PyTorch. Source code, pretrained weights, and step-by-step instructions for training and inference are available at https://github.com/AlexanderKroll/FoldVision.

### FoldVision model input

FoldVision voxelizes an atomic protein structure into a 3D grid with 1 Å resolution and five per-voxel channels (Figure 1). Each channel stores the local density of one of the five most common heavy-atom types: carbon, nitrogen, oxygen, sulfur, and phosphorus. For each voxel these values are stored in a 5-dimensional vector, and hence each protein is represented by a 4D array: three spatial dimensions plus one channel dimension.

Instead of one-hot encoding the presence of atoms at voxels, each atom contributes a continuous density to the 3D grid. For an atom of type *c* at position 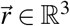, its contribution to the *c*-channel of the voxel centered at 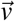 is exp 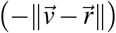. For each atom, we compute its contribution by iterating over all voxels within a distance of three voxels from the voxel containing the atom. This smooth representation reduces artifacts caused by discretizations and provides the network with local geometric context.

To limit empty space and memory consumption, proteins are fitted into the smallest of four cubic boxes 64^3^, 96^3^, 128^3^ and 160^3^ Å ^3^ that fully contains the protein. Structures that exceed the 160-voxel grid are cropped at the bounding box.

### Input data preprocessing

We obtained predicted structures in PDB format from the AlphaFoldDB [16]. To minimize data processing when running FoldVision, we converted every raw protein PDB file into a compact NumPy object before model training and validation.

We extract Cartesian coordinates for all atoms and we divide these coordinates by the target resolution (1 Å per voxel). The protein coordinates are then shifted so that all coordinates are non-negative and are at least four voxels away from the grid boundary. This ensures that the convolutional filters can adequately move over the protein surface.

Instead of storing a full 4D tensor for each protein, we save only non-zero entries as 5-tuples (*x, y, z, c, p*), where *x, y, z* are integer voxel indices, *c* ∈ {0,…, 4} is the channel (atom type), and *p* > 0 is the density value. We omit all voxel-channel combinations (*x, y, z, c*) with values of *p* = 0, resulting in much fewer coordinates than in a fully encoded 4D array. The resulting list is saved as a NumPy `.npy` file and that can be quickly converted into a 4D tensor for model input.

### Model architecture

FoldVision consists of a three-dimensional convolutional neural network (3D CNN) that operates directly on the atomic-density voxel grid described above. There are five convolutional blocks with cubic kernels sizes of 7, 5, 3, 3, and 3 voxels and feature depths of 128, 256, 512, 1 024 and 1 024. Within a block, one or two 3D convolutions are followed by group normalization and a ReLU activation. The spatial resolution is reduced by a filter stride of 2 in the first block and by max-pooling in all blocks (Table S1).

After the final block, the network yields a lower-resolution 3D grid whose voxels are represented by 1 024-dimensional feature vectors. Because the input box can vary (64–160 voxels per edge), the spatial size of this grid also varies. A global adaptive average-pooling layer compresses the grid to a fixed 1 024-dimensional protein embedding.

This protein embedding is fed into a prediction head consisting of a fully connected layer with 256 hidden units, batch normalization, ReLU activation, and 10 % dropout, followed by a linear output layer that returns either a scalar regression value or a probability, depending on the type of prediction task. For tasks involving a small molecule, a fixed-length small molecule embedding (see “Processing small molecule”) is concatenated with the protein embedding before the prediction head.

### Processing small molecules

For prediction tasks that involve a small molecule, every compound is first converted to its canonical SMILES string with explicit stereochemistry. We then use the MoLFormer-XL (MoLFormer-XL-both-10pct) [19], a SMILES Transformer pre-trained on ∼100 million molecules, to convert the SMILES strings into numerical embeddings: the model divides each SMILES string into distinct subparts, the so-called tokens, and computes for each token a numerical representation. The small molecule representations are kept fixed and we obtain a single embedding by averaging the token vectors element-wise.

### Rosetta energy term pre-training

We explored pre-training FoldVision as a multi-task regressor that predicts multiple Rosetta energy terms from a single input structure. For every Swiss-Prot protein in the training set we computed three groups of scores with PyRosetta (v3.13): (i) the full-atom score functions `ref2015`, (ii) the low-resolution centroid score functions `score3`, and (iii) four global descriptors from the `ProteinAnalyzer` class: total solvent accessible surface area, number of C_*α*_ –C_*α*_ contacts, count of hydrophobic residues, and the core-packing metric `two_core_each` (Table S2)

Additionally, if the pairwise Pearson correlation between two Rosetta score terms was greater than 0.95, one of these terms was randomly discarded, leaving 23 targets per protein. Each target was normalized to a zero mean and unit variance across the dataset. The network was then trained to minimise the mean-squared error averaged over these 23 outputs.

### Contrastive pre-training

We pre-trained FoldVision with a structure–sequence contrastive objective. For every Swiss-Prot entry we computed a fixed 1 280-dimensional sequence embedding with the pre-trained *ESM-2*_650M_ model [2]; these vectors remained frozen during training.

During pre-training, the 1 024-dimensional structure embedding produced by FoldVision is passed through a two-layer projection head to yield a 256-dimensional vector **z**^PV^. A similar projection head maps the ESM-2 embedding to **z**^ESM^ ∈ ℝ ^256^. For a mini-batch of *n* proteins we form the similarity matrix *S* with entries 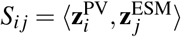 and optimize the symmetric InfoNCE loss with the temperature *τ* = 0.2 [40]. The loss encourages paired structure–sequence embeddings from the same protein to be close, and embeddings from different proteins to be distant.

### Model fine-tuning for protein encoders

We fine-tuned the 150 M- and 650 M-parameter versions of ESM-2 [2], ESM-IF [9], ProtT5 [3], ProstT5 [10], and 3DCNN-MQA [15] on every downstream task, using exactly the same train/validation/test splits as for FoldVision. We applied the same two-layer feed-forward prediction head (256 hidden units, batch normalization, ReLU activation, 10 % dropout, linear output) as for FoldVision to the updated protein embeddings. For small molecule dependent tasks this embedding is concatenated with the MolFormer-XL small molecule vector (see “Processing small molecules”). All model weights were updated during fine-tuning.

The models were optimized with the AdamW optimizer (weight decay 0.01); we screened learning rates between 1 × 10^−6^ and 1 × 10^−4^ on the validation set and selected the value that maximised the Matthews correlation coefficient for classification tasks and the coefficient of determination *R*^2^ for regression tasks. After training, we selected the model with the best validation performance, and we evaluated this model on the test set.

We used the esm-fair Python package to obtain the ESM models, the Hugging Face transformers library to obtain ProtT5 and ProstT5, and we re-implemented the 3DCNN-MQA model in PyTorch.

### Evaluation of GO-Term predictions

We evaluated Gene Ontology (GO) term prediction on the Molecular Function subset of the DeepGraphGO CAFA benchmark [41]. The corpus contains 35 092 proteins for training, 490 proteins for validation, and 426 proteins for testing, each annotated with experimentally verified GO terms.

Following the official CAFA protocol we report the *F*_max_ statistic, i.e., the maximum harmonic mean of precision and recall obtained when sweeping a threshold *τ* ∈ [0, 1] that binarises the predicted confidence scores *ŷ* ∈ [0, 1]. For a given *τ* we convert scores to binary labels 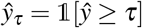 and compute precision and recall for every protein. Averaging these quantities over all *N* proteins yields

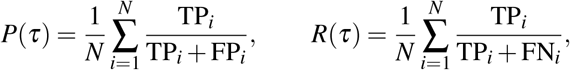

where TP_*i*_, FP_*i*_, and FN_*i*_ are the true-positive, false-positive, and false-negative counts for protein *i*. The *F*_1_ score at that threshold is

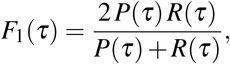

and *F*_max_ = max_*τ*_ *F*_1_(*τ*). The metric ranges from 0 (worst) to 1 (best) and is threshold-independent by design.

## Acknowledgements

We thank Deniz Sezer, Juan Farina, Yvan Rousset, and Julia Kroll for helpful discussions. Computational support and infrastructure were provided by the “Centre for Information and Media Technology” (ZIM) at the University of Düsseldorf (Germany). This work was funded through grants to MJL by the European Union (ERC AdG “MechSys”–Project ID 101055141) and by the Deutsche Forschungsgemeinschaft (DFG, German Research Foundation: CRC 1310, and, under Germany’s Excellence Strategy, EXC 2048/1–Project ID 390686111).

## Supplementary Information

### Supporting Tables S1-S3

**Supplementary Table S1.** Layer–by–layer configuration of the FoldVision model with 5 convolutional blocks.

| # | Layer | In Ch. | Out Ch. | Kernel / Pool | Stride |
| --- | --- | --- | --- | --- | --- |
| 1 | Conv3d | 5 | 128 | $7 \times 7 \times 7$ | 2 |
| 2 | GroupNorm3d+ReLU | 128 | 128 | – | – |
| 3 | MaxPool3d | – | – | $2 \times 2 \times 2$ | 2 |
| 4 | Conv3d | 128 | 256 | $5 \times 5 \times 5$ | 1 |
| 5 | GroupNorm3d+ReLU | 256 | 256 | – | – |
| 6 | Conv3d | 256 | 256 | $5 \times 5 \times 5$ | 1 |
| 7 | GroupNorm3d+ReLU | 256 | 256 | – | – |
| 8 | MaxPool3d | – | – | $2 \times 2 \times 2$ | 2 |
| 9 | Conv3d | 256 | 512 | $3 \times 3 \times 3$ | 1 |
| 10 | GroupNorm3d+ReLU | 512 | 512 | – | – |
| 11 | Conv3d | 512 | 512 | $3 \times 3 \times 3$ | 1 |
| 12 | GroupNorm3d+ReLU | 512 | 512 | – | – |
| 13 | MaxPool3d | – | – | $2 \times 2 \times 2$ | 2 |
| 14 | Conv3d | 512 | 1024 | $3 \times 3 \times 3$ | 1 |
| 15 | GroupNorm3d+ReLU | 1024 | 1024 | – | – |
| 16 | Conv3d | 1024 | 1024 | $3 \times 3 \times 3$ | 1 |
| 17 | GroupNorm3d+ReLU | 1024 | 1024 | – | – |
| 18 | MaxPool3d | – | – | $2 \times 2 \times 2$ | 2 |
| 19 | Conv3d | 1024 | 1024 | $3 \times 3 \times 3$ | 1 |
| 20 | GroupNorm3d+ReLU | 1024 | 1024 | – | – |
| 21 | Conv3d | 1024 | 1024 | $3 \times 3 \times 3$ | 1 |
| 22 | GroupNorm3d+ReLU | 1024 | 1024 | – | – |
| 23 | MaxPool3d | – | – | $2 \times 2 \times 2$ | 2 |

**Supplementary Table S2.** Rosetta energy terms and global descriptors used as regression targets during pre-training.

| Score terms |  |
| --- | --- |
| cenpack | omega |
| contact_all | p_aa_pp |
| dslf_fa13 | pair |
| env | pro_close |
| fa_atr | rama_prepro |
| fa_elec | ref |
| fa_sol | rg |
| hbond_bb_sc | total_hydrophobic |
| hbond_lr_bb | total_sasa |
| hbond_sc | total_score |
| hbond_sr_bb | two_core_each |
| lk_ball_wtd | vdw |

**Supplementary Table S3.** Dataset sizes for the four benchmark tasks used in this study.

| Dataset | Training size | Validation size | Test size |
| --- | --- | --- | --- |
| Enzyme–Substrate | 733 139 | 5 282 | 13 086 |
| Transporter–Substrate | 23 858 | 2 628 | 6 014 |
| GO Terms (Molecular Function) | 35 092 | 490 | 426 |
| IC<sub>50</sub> (BindingDB) | 932 442 | 50 000 | 51 720 |
| Davis (cold target) | 23 868 | 3 094 | 3 094 |

